# Disordered phasic relationships between hippocampal place cells, theta, and gamma rhythms in the Ts65Dn mouse model of Down Syndrome

**DOI:** 10.1101/2020.09.17.301432

**Authors:** H.C. Heller, A. Freeburn, D.P. Finn, R.G.K Munn

## Abstract

Down Syndrome (DS) in humans is caused by trisomy of chromosome 21 and is marked by prominent difficulties in learning and memory. Decades of research have demonstrated that the hippocampus is a key structure in learning and memory, and recent work with mouse models of DS have shown changes in spectral coherence in the field potentials of hippocampus and regions important for executive function such as prefrontal cortex. One of the primary functional differences in DS is thought to be an excess of GABAergic innervation from Medial Septum (MS) to regions such as hippocampus. In these experiments, we probe in detail the activity of region CA1 of the hippocampus using *in vivo* electrophysiology in the Ts65Dn mouse model of DS in comparison to their non-trisomic 2N littermates. We find changes in hippocampal phenomenology that suggest that MS output, which drives theta rhythm in the hippocampus, is strongly altered. Moreover, we find that this change affects the phasic relationship of both CA1 place cells and gamma rhythms to theta. Since the phasic relationship of both of these aspects of hippocampal phenomenology to theta are thought to be critical for the segregation of encoding and retrieval epochs within hippocampus, it is likely that these changes are the neural substrates of the learning and memory deficits seen both in human DS and animal models such as Ts65Dn.

## Introduction

The human chromosomal disorder Down Syndrome (DS) involves partial or complete trisomy of chromosome 21. DS produces a spectrum of interrelated morphological changes, including skeletal changes^1,2^, cardiac abnormalities^3–5^, as well as deficits in executive function^6,7^. Individuals with DS exhibit generalized deficits in learning and memory^8,9^, including specific deficits reminiscent of hippocampal dysfunction^10^ Several mouse models of DS exist, with varying amounts of overlap in trisomic genes between murine and human. The Ts65Dn mouse model used in the present study exhibits partial trisomy of murine chromosome 16, which contains many of the candidate genes for the changes seen in human DS. These mice exhibit many of the changes seen in human DS, such as skeletal and cardiac abnormalities as well as problems with learning and memory.

Although the precise mechanism of memory disorder in Ts65Dn animals is not known, these animals show changes in output from medial septum (MS) to hippocampus^11,12^, which is thought to be the key generator of local field potential (LFP) theta^13–17^, which in turn is key to normal hippocampal function^18–26^. Specifically, Ts65Dn animals show an altered excitatory/inhibitory balance in hippocampus, with marked decreases in cholinergic^11,12^, and substantially greater GABAergic signalling^27,28^. A likely substrate for these changes within the hippocampus is the medial septum, which sends large cholinergic and GABAergic projections to hippocampus^13,29–31^. Individual place cells in area CA1 of the hippocampus display robust precession of their firing through the local theta wave as animals traverse the region of space that comprises the “place field” for each cell^32–35^. The phasic relationships among the different aspects of hippocampal phenomenology are central to normal hippocampal function. Segregation of single unit activity on the theta wave is thought to route information flow to support either encoding or retrieval^22,36^. The switching of LFP gamma into “fast” and “slow” modes switches hippocampal place cells into prospective and retrospective coding modes^37^. The two different speeds of gamma themselves are also segregated on the theta cycle, and entrain CA1 with either CA3 (slow gamma) or MEC (fast gamma). Slow gamma occurs predominantly on the falling portion of the cycle, and fast gamma occurs mainly near the trough^38^. This spectral separation is mirrored by spatial separation in laminar depth, with slow gamma from CA3 arriving mainly proximal to the soma of CA1 pyramidal cells in stratum radiatum, and faster gamma from layer III of MEC typically arriving in the distal portion of the apical dendrites of CA1 pyramidal cells in stratum lacunosum moleculare^39^. The fact that the different frequencies of gamma oscillations are associated with switching place cells in CA1 from retrospective to prospective coding underlines the importance of gamma rhythms in segregation of information flow^37^.

Other spectral components of hippocampal activity are also key to normal function. Sharp wave ripples within the hippocampus have also been shown to be critical in the consolidation of memory; in awake animals, place cells rapidly preplay firing patterns in an extremely temporally compressed form that is predictive of their future movement^40,41^. Closed-loop experiments preventing the generation of hippocampal sharp-wave events through electrical stimulation^42^ clearly show the necessity of these events in the consolidation of spatial memory. Increased duration of sharp-wave ripples has been shown to be correlated with increases in memory demand, and artificially increasing the duration of ripples optogenetically results in improved spatial memory^43^

Output from medial septum is abnormal in Ts65Dn animals, and recent evidence has shown excessive GABAergic innervation of the dentate gyrus in these animals^27^. Moreover, GABA antagonism restores learning and memory in Ts65Dn mice^44–46^. This constellation of evidence leads to the hypothesis that since input to hippocampus from MS is dysfunctional in DS, hippocampal function in TS animals may be abnormal in ways that parallel the changes in human DS that result in deficits in learning and memory. Recent experiments using mouse models of DS strongly suggest spectral phase abnormalities between the hippocampus and prefrontal cortex, and differences in phase-amplitude coupling between theta and gamma in the hippocampus^47^. In these experiments, we use tetrode-based single-unit electrophysiology to record place cells within CA1 of the hippocampus, co-record hippocampal LFP, and examine changes in spectral coherence within the hippocampus that may underlie the central learning and memory deficits in DS. Given the altered input from septum to hippocampus in Ts65Dn animals, and the learning and memory deficits associated with this model, we hypothesise that there will be changes in the phasic relationships of activity in the hippocampus that explain the behavioral outcomes. We therefore carefully examine the activity of single units in hippocampus with respect to the ongoing rhythmic oscillations in hippocampus, and the phasic relationships of the different frequency bands within hippocampus to each other. Changes in these aspects of hippocampal phenomenology will have critical implications for the ability of the system to normally function.

## Results

Single units recorded from Ts65Dn animals and their non-trisomic 2N littermate controls were isolated as animals explored a 40 x 40 cm black polycarbonate arena. A total of 179 neurons were isolated in the Ts65Dn group, and 72 from the 2N Control group. These neurons were manually separated into putative interneurons and pyramidal cells on the basis of their location specificity, firing rate, and waveform. After removing all interneurons, there were n = 119 neurons remaining in the Ts65Dn group and n = 64 neurons remaining in the 2N control group.

### The spatial properties of Ts65Dn place cells are normal

Non-interneuron cells recorded from the hippocampus of 2N and Ts65Dn animals were classified as place-encoding if they passed the thresholds for the standard place-cell metrics information content, sparsity, and selectivity generated by the temporal shuffling of spikes (see methods). In addition, cells were included only if they had identified fields covering not less than 2% of the environment and not more than 50% of the environment in total. (see methods). This left n = 107 cells in the Ts65Dn group and n = 42 cells in the 2N control group. Place cells in both 2N and Ts65Dn groups appeared similar (Figure 1A). There was no difference in the spatial information content (Figure 1B, Mean information content ± SEM (bits/spike), Ts65Dn = 0.269 ± 0.033; 2N Control = 0.258 ± 0.065, Z = 1.323, p = 0.186, n.s), spatial sparsity (Figure 1C, mean sparsity ± SEM, Ts65Dn = 0.744 ± 0.017; 2N Control = 0.772 ± 0.028, Z = 1.314, p = 0.189, n.s), or ratio of infield to outfield firing rates between 2N and Ts65Dn cells (Figure 1D, mean infield/outfield firing rate ratio ± SEM, Ts65Dn = 3.03 ± 0.16, 2N Control = 3.03 ± 0.35, Z = 0.437, p = 0.662, n.s). The size of the place fields was similar between the groups of cells (mean % place field coverage ± SEM; Ts65DN = 14.37 ± 1.09, 2N Control = 12.78 ± 1.65, Z = 1.07, p = 0.287, n.s). However, Ts65Dn place cells fired significantly fewer spikes in bursts of spikes than control (mean spikes in bursts/single spikes ± SEM, Ts65Dn = 0.581 ± 0.116, 2N Control = 0.612 ± 0.102, Z = 2.44, p = 0.0146).

**Figure 1.**
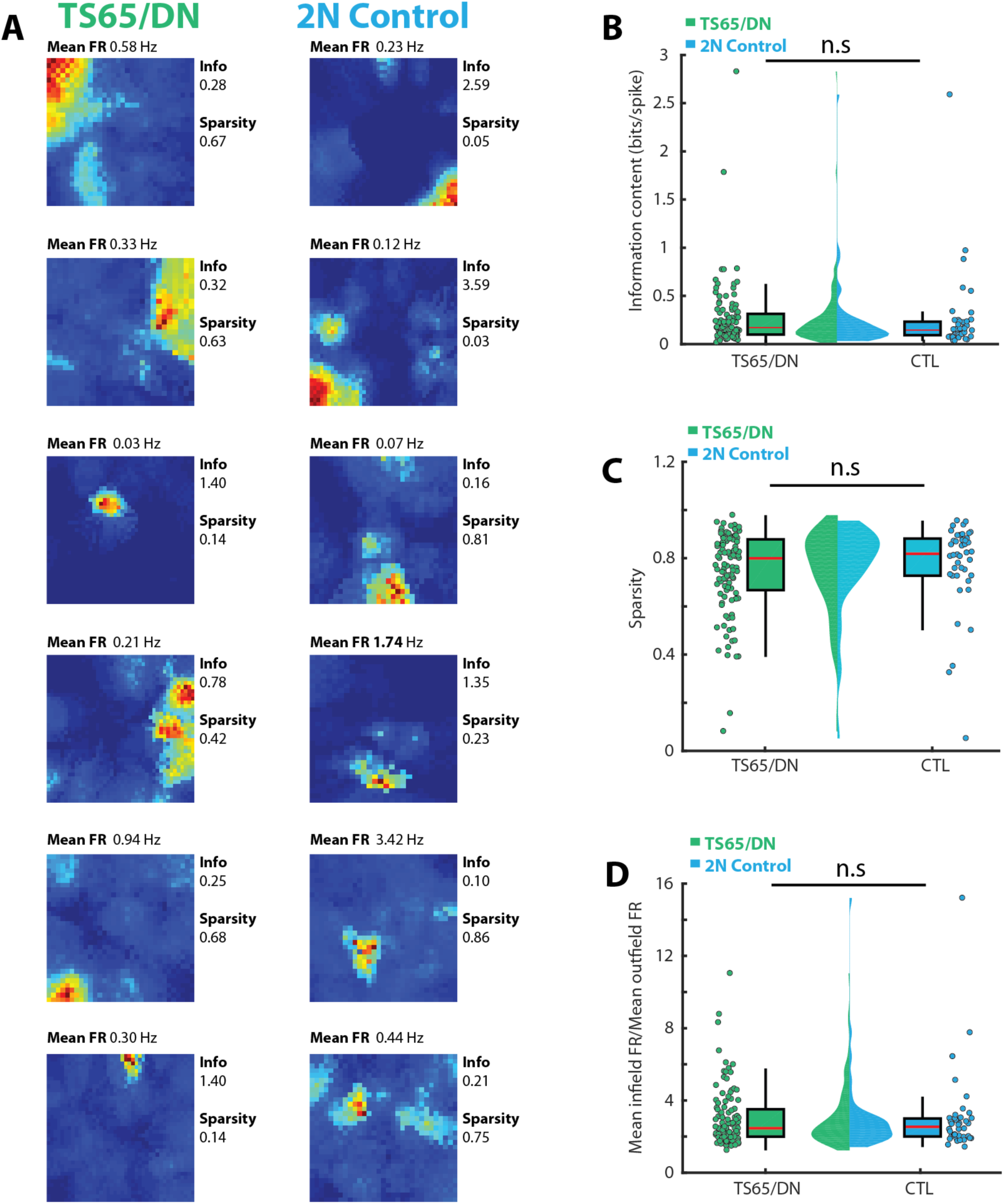
The spatial properties of Ts65Dn place cells are normal. **A)** 12 color-coded rate maps of place cells recorded from Ts65Dn (left column) and 2N Control littermate mice (right column). Place cells from both groups appeared normal, with well-defined firing fields. “hotter” colors represent higher firing rates. Beside each cell is its spatial information content and spatial sparsity. The mean firing rate of each cell is shown above each. **B)** The information content relayed by each spike (in bits) was similar between Ts65Dn and 2N control. **C**) The spatial sparsity of the spikes fired by place cells in each group was similar, indicating relatively well-defined place fields in both groups of animals. **D**) The ratio of spikes fired in identified fields to spikes fired outside identified fields was similar between place cells recorded from Ts65Dn and 2N Control mice. n.s = not significant.

### The frequency of hippocampal theta rhythm is higher in Ts65Dn animals than 2N Control

Sessions from which single units were recorded were used for the analysis of hippocampal theta rhythm (Ts65Dn, sessions = 21; 2N Control, sessions = 13). Both Ts65Dn and 2N Control animals displayed a typical dominant theta rhythm in their hippocampal LFP (Figure 2A). The mean frequency of theta rhythm in Ts65Dn animals was significantly higher than 2N control (Figure 2A,D. Mean frequency (Hz) ± SEM, Ts65Dn = 8.42 ± 0.029 Hz, 2N Control = 8.25 ± 0.040 Hz, Z = 3.083, p = 0.002). While theta frequency is linearly correlated with running speed^48^, The difference in frequency between Ts65Dn and 2N control theta frequency was not due to difference in running speed; although the Ts65Dn animals ran faster on average than 2N Control (mean speed ± SEM (cm/s); Ts65Dn = 14.21 ± 0.614 cm/s, 2N Control = 8.96 ± 0.355 cm/s, Z = 4.64, p = 3.44^-6^) the intercept of a linear regression through the speed/frequency data was still greater for Ts65Dn animals compared to control (Figure 2B, Theta intercept ± SEM (Hz), Ts65Dn = 8.349 ± 0.040 Hz, 2N Control = 8.115 ± 0.067 Hz, Z = 2.693, p = 0.0071). The linear relationship of theta frequency to running speed appeared to be flatter in Ts65Dn animals than control, but this apparent difference did not reach significance (mean slope ± SEM (Hz/cm/s), Ts65Dn = 0.0063 ± 0.0012, 2N Control = 0.0084 ± 0.0029, Z = 0.390, p = 0.697, n.s). While there was no overall difference in the mean normalized power over the entire theta band between Ts65Dn and 2N control animals (Figure 2D) this was due to a rightshifted preferred band of theta at higher frequencies in the Ts65Dn animals. The peak power of theta was higher in Ts65Dn animals than 2N control (Figure 2D, normalized peak theta power ± SEM; Ts65Dn = 0.012 ± 7.66^-4^, 2N control = 0.009 ± 5.16^-4^, t(1,33) = 2.464, p = 0.0193).

**Figure 2.**
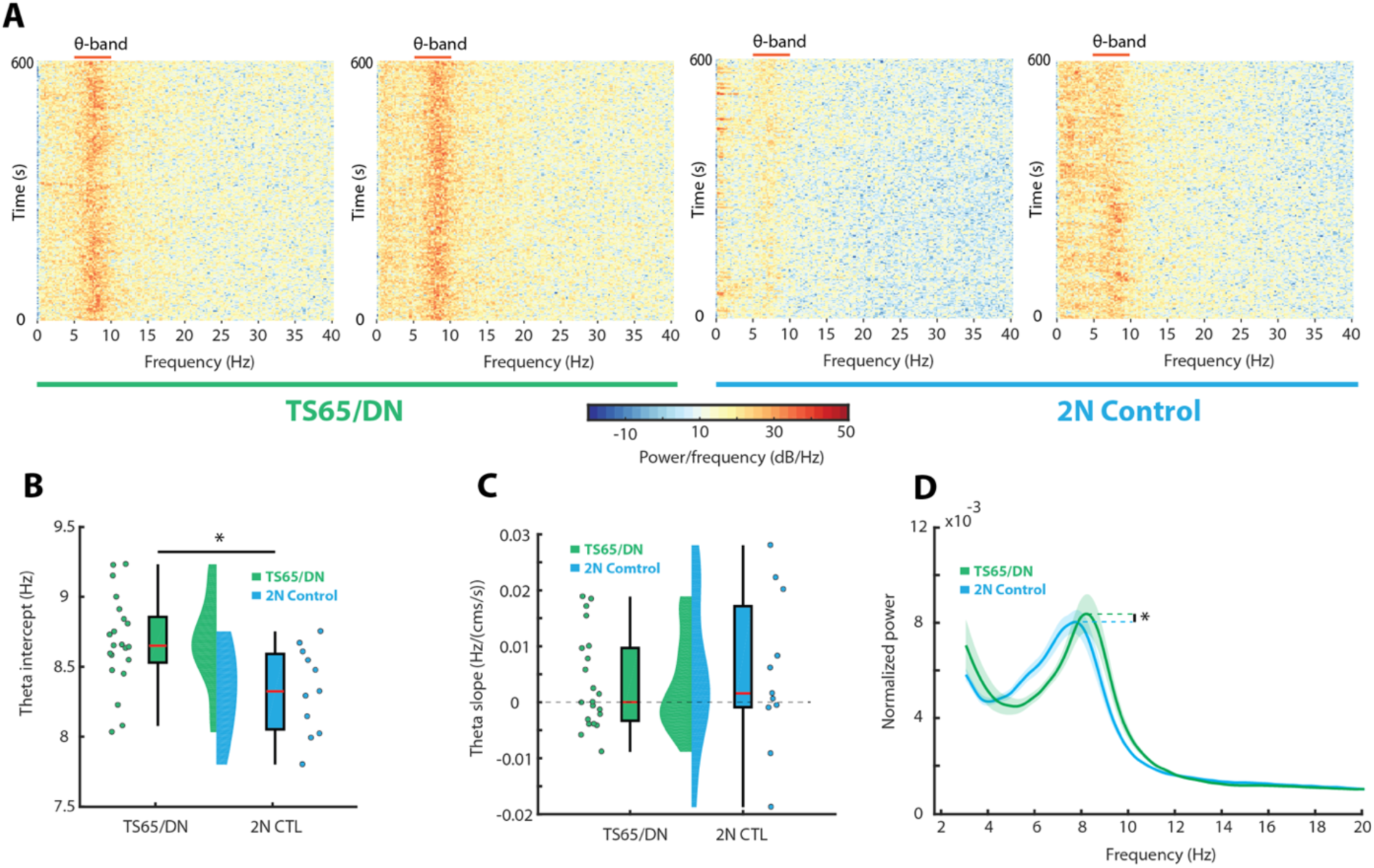
The frequency of hippocampal LFP theta rhythm is higher in Ts65Dn animals than 2N control. **A)** Spectrograms showing the spectral power between 0-40Hz over four individual 600-second recording sessions (two Ts65Dn, left, green; two 2N control, blue, right). “hotter” colors indicate higher power. The approximate location of the theta band (5-11Hz) is illustrated for each spectrogram with a red bar above the band. A strong band of high power in the theta range is evident in each recording; typical for recordings from hippocampus. **B)** The intercept generated by a linear regression through the running speed/frequency data generated from each recording session for Ts65Dn (green) and 2N (blue) animals. Boxplots for each group show the interquartile range (box extents), and median values (red line) for each group. Colored back-to-back half-violin plots show the relative density of values for each group. **C)** The slope of the linear regression through the running speed/frequency data for Ts65Dn (green) and 2N control (blue) recording sessions. A black dotted line illustrates 0 slope. Other features of the figure are as in (B). **D)** Normalized spectral power in recordings made from Ts65Dn (green) and 2N control (blue) animals in the 3-20Hz band. Mean power is illustrated by a solid line, while the shaded region illustrates SEM. Coloured dashed lines illustrate the mean power at the peak of theta in each group. *p < 0.05

### Ts65Dn hippocampal cells are more likely to be theta rhythmic, and are highly phase locked to hippocampal theta rhythm compared to 2N Control

Although there was no difference in the spatial characteristics of place cells in Ts65Dn animals compared to control, cells in these animals were much more tightly phase locked to the ongoing LFP theta than control (Figure 3A,C,D). The phasic directional vector was much larger in Ts65Dn cells (n = 119) than 2N Control (n = 63, Mean vector length ± SEM; Ts65Dn = 0.143 ± 0.007, 2N Control = 0.083 ± 0.009, Z = 5.054, p = 4.319^-7^). Demonstrating that spikes most often occurred on regular phases of theta in Ts65Dn cells. In contrast, the firing of 2N control cells was distributed normally through phases of theta as the animal moved through the firing field of each cell. Ts65Dn cells tended to lock to the peak or trough of theta (Figure 3B,D,E,F); the direction of the mean vector of these cells was significantly clustered compared to 2N control (F(1,180) = 38.75, p = 3.3^-9^). The mean phase direction of 2N control cells was evenly distributed at all phases of theta (U = 146.2, p = 0.100). In contrast, the phase directions of the directional vector of Ts65Dn cells were highly clustered (U = 160.1, p = 0.001). Using a method for estimating phase precession in two dimensions by isolating individual passes through a firing field^49^ we discovered relatively poor strength of a circular-linear regression of the pass index with theta phase in both groups, with no significant difference in correlation between them (Mean r ± SEM, Ts65Dn = −0.03 ± 0.003; 2N Control = −0.031 ± 0.003, Z = 0.61, p = 0.54). In agreement with the relatively strong phase locking of Ts65Dn neurons to the peak of theta, we find that the intercept of the linear-circular regression in the Ts65Dn animals was distributed differently from 2N Control animals, and was weighted toward the peak of theta (Figure 3G-H, mean intercept ± SEM, Ts65Dn = 214.02 ± 10.39°, 2N Control = 188.8 ± 14.13°, KS = 0.213, p = 0.038).

**Figure 3.**
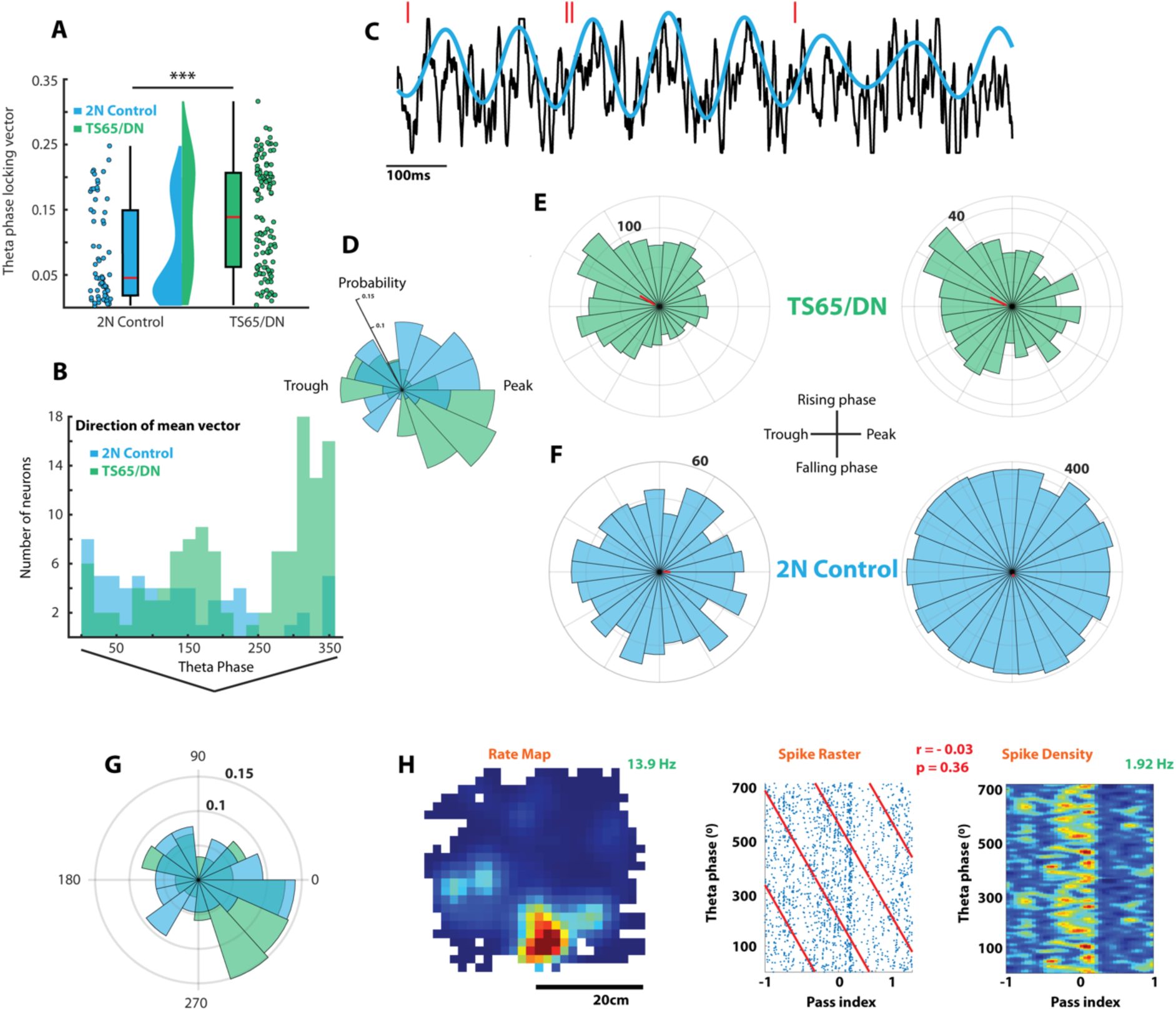
Ts65Dn place cells are abnormally phase locked to hippocampal theta. The mean directional vector generated from the distribution of the phase angle at which the spikes of each cell recorded from Ts65Dn (green) and 2N control (blue) animals. Boxplots show the interquartile range (boxes), and median (red line) of the distributions, while half-violin plots show the density of values in each distribution. **B)** Histogram showing the number of cells with a particular mean direction vector (as in (A)) from each cell occurred in the Ts65Dn (green) and 2N control (blue). **C)** Example of a raw hippocampal LFP trace (black) and a bandpass of this signal (blue), along with the co-occuring spikes of a co-recorded single unit (red lines). **D)** Polar histogram showing the probability that a given cell in the Ts65Dn (green) and 2N Control (blue) groups have a particular mean directional vector. **E)** Polar histograms showing the number of spikes fired at specific phases of theta by two example Ts65Dn cells (green). Bold numerals indicate the number of spikes in the outermost ring they touch, while red lines illustrate the magnitude (by length) and direction of the mean vector produced from these distributions. **F)** As in E, but for cells recorded from 2N Control animals. A place cell (left) and interneuron (right) are shown. G) Polar histogram showing the probability that a circular-linear regression of pass location and theta phase has a particular intercept for Ts65Dn (green) and 2N Control (blue) place cells. Bold numbers indicate the probability represented by the ring to which they are attached. The cardinal directions of the plot indicate the theta phase from 0/360 (peak), 90 (falling phase), 180 (trough), and 270 (rising phase). **H)** Pass analysis of an example place cell. The leftmost plot shows the rate map of the cell, while the green number shows the peak infield firing rate. The middle plot shows the location of the spikes in pass and phase space. The pass index ranges from −1 (maximally before field center), 0 (field center) and 1 (maximally after entry to field). The red lines show the linear-circular intercept, and the red numbers the rho and p-value of the fit. The rightmost plot shows the binned rate density of the pass/phase raster plot (hotter colors: more spikes). The green number is the firing rate indicated by the hottest colour. ***p < 0.001

Determination of the intrinsic rhythmicity of the spiking of cells in both groups was carried out using the Maximum Likelihood Estimation (MLE) method as in Climer (2015)^50^. This method derives a rhythmic modulation index ranging from 0 (no rhythmicity) to 1 (completely rhythmic). Cells recorded from Ts65Dn animals had significantly higher rhythmicity indices than cells recorded from 2N Control animals (Figure 4 A-C) Rhythmicity index ± SEM; Ts65Dn = 0.487 ± 0.025, 2N Control = 0.320 ± 0.037, Z = 3.83, p = 1.26^-4^). These data indicate a much greater degree of intrinsic theta rhythmicity in the cells of Ts65Dn animals. The MLE method also determines whether the rhythmic fit is a significantly better fit to the spiking data than a flat, non-rhythmic fit. Ts65Dn cells were much more likely to be better fit by a rhythmic fit than a flat decay than 2N Control (Figure 4D, percent significantly rhythmic; Ts65Dn = 56.94 %, 2N Control = 26.15 %, χ^2^ = 12.27, p = 0.0005).

**Figure 4.**
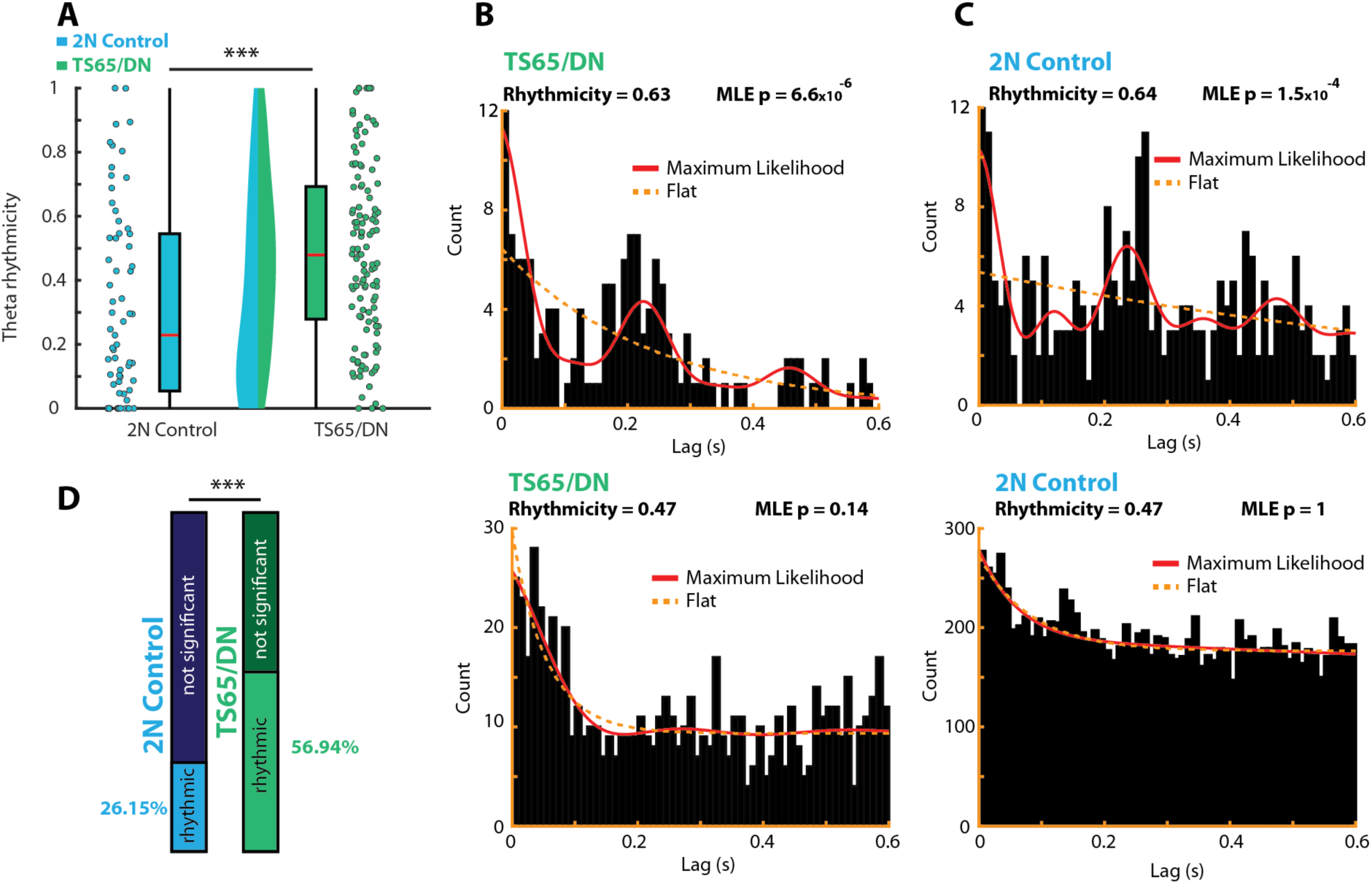
Ts65Dn place cells are intrinsically more theta rhythmic than 2N control. **A)** The theta rhythmicity of the spikes fired by each cell (0, non-rhythmic, 1, maximally rhythmic) in the Ts65Dn (green) and 2N control (blue) groups. Boxplots show the median (red lines) and interquartile range (box extents), while half-violins illustrate the relative distribution of value density. **B)** Example autocorrelograms of a theta rhythmic (top) and non-rhythmic (bottom) cell recorded from Ts65Dn animals. The red solid line illustrates the maximum likelihood fit, while the orange dotted line shows the fit with no rhythmicity, as in Climer et.al (2015). The probability that the maximum likelihood fit is not a better than a flat, non-rhythmic fit is shown in the top right of each histogram. **C)** as in **B)**, but for a rhythmic cell (top) and a non-rhythmic cell (bottom) recorded from 2N Control animals. **D)** The proportion of cells recorded from the 2N Control (blue bar) and Ts65Dn (green bar) groups that are better fit by a rhythmic fit (light colors) compared to the proportion of cells better fit by a flat, non-rhythmic fit (darker colors). Cells from the Ts65Dn group were more often rhythmic than cells from the 2N Control group. *** p < 0.001.

### Increased coherence and abnormal phasic relationships between theta and gamma rhythms in Ts65Dn animals

Spectral coherence between theta and gamma bands (Figure 5A,B) overall was greater in the Ts65Dn animals than the 2N controls (Figure 5C-F mean theta/gamma coherence ± SEM; Ts65Dn = 0.032 ± 0.007, 2N control = 0.016 ± 0.002, t(1,32) = 1.71, p = 0.049) The peak of coherence between theta and fast gamma appeared to occur at higher frequencies in Ts65Dn animals than control (Figure 5C, peak coherence ± SEM; Ts65Dn = 75.52 ± 2.77Hz, 2N Control = 71.54 ± 0.79Hz). Subtraction of the mean cross-frequency coherence in the theta band of Ts65Dn animals from 2N control demonstrated greater coherence between both fast and slow gamma and higher frequencies of theta in the Ts65Dn animals (Figure 5D), however, when the coherence for each recording session was normalized within each session, it became apparent that there was relatively greater coherence between both gamma bands and lower frequency theta in 2N control animals (Figure 5D).

**Figure 5.**
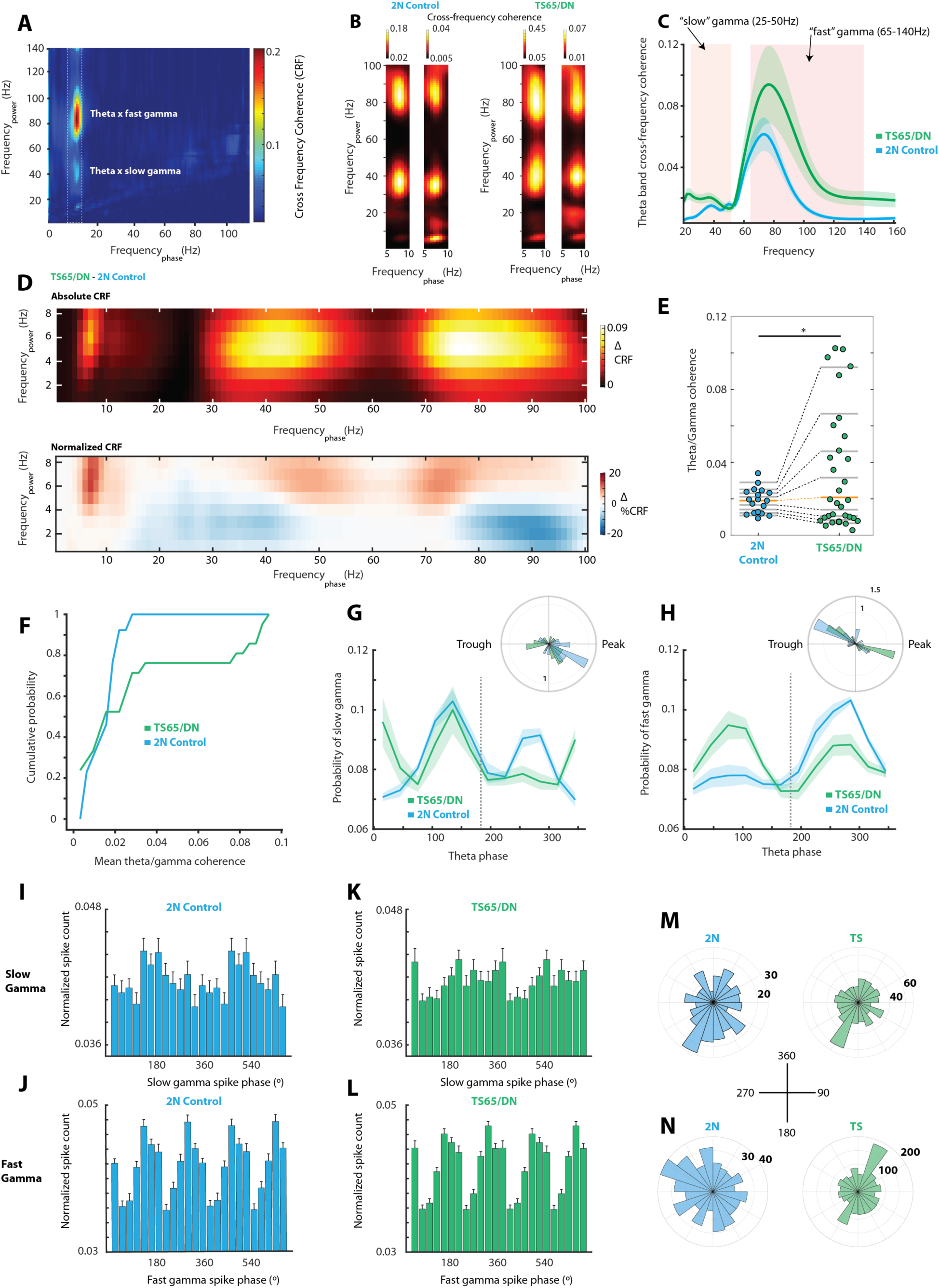
Coherence between hippocampal theta and gamma rhythms is altered in Ts65Dn mice. **A)** Example cross-frequency coherence spectrograph showing the coherence between frequency bands. In this example, spectral energy in the theta band is highly coherent with energy in the fast gamma band, and there is moderate coherence between theta and slow gamma. A white dashed rectangle illustrates the location of the theta phasic band. **B)** Cross-frequency coherence spectrograms for two 2N control animals (left pair) and two Ts65Dn animals (right pair). The cross-frequency spectrograms are restricted to the theta band for clarity. All four animals show strong coherence between theta and both gamma bands. **C)** The mean cross-spectral coherence (solid lines) ± SEM (shaded regions) between theta and other frequencies. The 2N control group (blue) shows strong spectral coherence between theta and both slow and fast gamma (shaded regions), while the cross-frequency coherence in the theta band is larger in both the slow and fast gamma bands in the Ts65Dn group (green), and appears to peak at higher frequencies of fast gamma. **D)** Top; the mean cross-frequency coherence over all recordings in the 2N control group subtracted from the mean cross-frequency coherence over all recordings in the Ts65Dn group. This plot shows the relatively higher theta/gamma coherence in the Ts65Dn animals. Bottom: As in the top plot, but each cross-frequency coherence matrix for each recording was internally normalized such that the maximum value in each was 1. This plot demonstrates the relatively greater cross-spectral coherence between both bands of gamma and lower theta frequencies in the 2N control animals, while the greatest cross-frequency coherence in Ts65Dn animals occurred at higher frequencies of theta and slow gamma, and lower frequencies of fast gamma. **E)** Quantile plot showing the mean coherence between theta and both gamma bands for 2N control (blue) and Ts65Dn (green) animals. Each point is the mean coherence in one recording session. The median values of each group are shown with orange line, while the upper and lower quartiles of each group are shown with grey lines. The corresponding quartiles are joined with dashed lines. **F)** The cumulative probability that a recording has a given mean coherence between theta and gamma. Ts65Dn (green line) recordings were more likely to have greater coherence between theta and gamma than 2N control (blue line). **G)** The mean probability (solid lines) ± SEM (shaded area) of slow gamma occurring over phases of the theta cycle for 2N (blue) and Ts65Dn (green) recording sessions. The dashed grey line indicates the peak of theta (180°). Inset polar histogram shows the relative probability for all instances of slow gamma that slow gamma occurs at a particular phase of theta (blue bars, 2N Control; green bars, Ts65Dn) **H)** As in **(G)**, but for the probability that fast gamma occurs at various phases of theta (main figure) and relative probability of each fast gamma epoch occurring at a particular phase of theta (inset polar histogram). **I**,**J)** The mean normalised spiked counts ± SEM at each phase of slow (I) and fast (J) gamma for 2N Control cells. **K**,**L)** As in (**I**,**J)** but for cells recorded from Ts65Dn animals. **M)** Polar histograms showing the number of spikes fired at a specific phase of slow gamma for a cell recorded from a 2N Control animal (blue, left) and a Ts65Dn animal (green, right). Inset numbers denote numbers of spikes in each ring. **N)** as in **(M)**, but for two different cells and their phasic relationship to fast gamma. Inset cross shows the phasic direction on the gamma wave. *p < 0.05

Previous work has shown a phasic relationship between the theta wave and the different bands of gamma, with gamma typically occurring on appearing to be superimposed on, theta oscillations Also, fast and slow gamma typically do not appear on the same theta cycle. We find that this is true of both 2N control (mean correlation) and Ts65Dn (mean correlation) animals. The location of the superposition of fast and slow gamma on theta is also typically segregated, with fast and slow gamma preferring different phases of theta. In the 2N control animals, this phasic segregation is apparent, as in Colgin et.al (2009)^38^. Slow gamma typically occurs close to the peak of theta (Figure 5G, Rayleigh test, Z = 6.59, p = 8.057^-4^), while fast gamma occurs on the falling phase of theta nearer to the trough (Rayleigh test, Z = 3.99, p = 0.016; Figure 5H). However, in Ts65Dn animals, there is markedly less phasic separation. While the mean theta phase of slow gamma in 2N animals is unipolar and relatively specific to the rising phase of theta The mean theta phase of Ts65Dn slow gamma is relatively evenly distributed (Figure 5G, Rayleigh test, Z = 0.69, p = 0.505, n.s.), as is the mean theta phase of fast gamma (Figure 5H, Rayleigh test, Z = 0.135, p = 0.877, n.s.). While the mean theta phase of slow gamma was not different between 2N control and Ts65Dn animals (Watson-williams test, F(1,36) = 3.23, p = 0.081, n.s), Fast gamma in Ts65Dn animals often occurred at opposite phases of theta than 2N control (Watson-Williams test, F(1,36) = 4.34, p = 0.044).

Previous research has demonstrated that neurons within the hippocampus are sometimes phase locked to gamma oscillations ^37,38,51^. In both 2N and Ts65Dn animals, we find that, on average, cells are phasically entrained to both slow and fast gamma (Figure 5I-N). However, spikes fired by neurons in 2N control animals were equally likely to be phase locked to slow and fast gamma (14/61 phase locked to slow gamma, 10/61 phase locked to fast gamma, Q = 0.830, p = 0.36) while Ts65Dn neurons were more likely to be phase locked to slow gamma compared to fast (13/114 fast, 25/114 slow, Q = 4.55, p = 0.033)

### Sharp-wave ripple duration is increased in Ts65Dn animals compared to 2N control

While there were prominent changes in both spike and gamma entrainment to the ongoing LFP theta rhythm in the Ts65Dn animals compared to control, there were no clear differences in the appearance of sharp-wave ripples or the phasic relationships of spikes to ripples between Ts65Dn animals and control. Ripples occurred approximately as often in the LFP of Ts65Dn animals as control (Figure 6A,E, mean ripple frequency ± SEM (Hz), Ts65Dn = 0.280 ± 0.031; 2N Control = 0.343 ± 0.055, Z = 1.028, p = 0.304, n.s). Spikes in both groups occurred less than a quarter of the time during ripples (Figure 6D, mean percentage spikes fired during ripples ± SEM, Ts65Dn = 12.30 ± 0.504 %; 2N Control = 11.32 ± 0.696 %, Z = 0.946, p = 0.329, n.s). While ripples occurred approximately as often in the LFP of Ts65Dn animals as 2N control animals, ripple episodes were each of much longer duration in Ts65Dn animals (Figure 6B,C, mean ripple duration ± SEM (msec), Ts65Dn = 60.7 ± 3.1 msec; 2N Control = 21.7 ± 0.09 msec, Z = 9.406, p = 5.17^-21^). Ts65Dn ripple duration was also much more variable than 2N Control ripples, which were usually approximately the same length (Levene’s test = 233.63, p < 0.0001).

**Figure 6.**
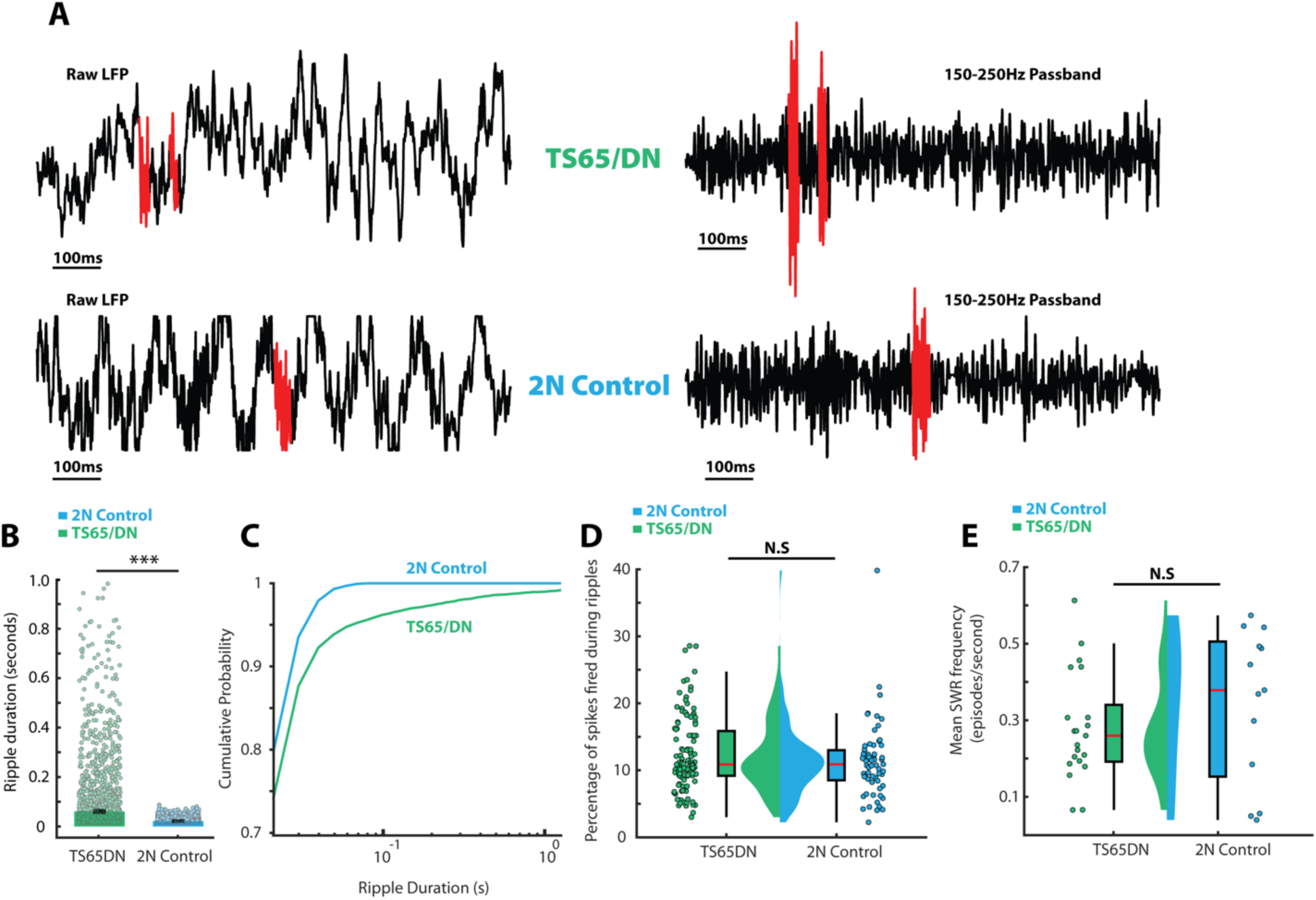
The duration of hippocampal sharp-wave ripples, but not their incidence, was higher in Ts65Dn animals than 2N control. **A)** Example traces of local field potential recordings from Ts65Dn (green, top row) and 2N control (blue, bottom row). Raw LFP traces are shown on the left, and the same trace filtered with a 150-250Hz passband filter are shown on the right. Sharp-wave ripples are shown in red. **B)** The duration of each sharp-wave ripple recorded from Ts65Dn (green) and 2N control (blue) animals. Solid bars illustrate the mean values, and black bars show the standard error. **C)** The percentage of spikes fired by each cell recorded from Ts65Dn animals (green) and 2N controls (blue) that occurred during a sharp-wave ripple in the concurrently recorded local field potential. Boxplots illustrate the interquartile range, while the median for each group is shown on the boxplot as a red line. Half violin plots show the density of the distribution of values in each group. **D)** The mean frequency with which sharp wave ripples occurred (in number of ripples per second) in the local field potential of each recording from Ts65Dn (green) and 2N control (blue) animals. Plot features are as in **(C)**. *** p < 0.001, n.s = not significant

## Discussion

Theta rhythm in the hippocampus has been shown extensively to be critical in the formation of episodic memory and for successful spatial navigation. Specifically, the phasic relationship of theta to other aspects of hippocampal activity such as gamma is critically important in the correct routing or combination of the disparate streams of information converging on the hippocampus from regions such as entorhinal and frontal cortex. The Ts65Dn mouse model of DS is known to exhibit the learning and memory deficits central to the syndrome^52^, but in these experiments we paradoxically observe what initially appears to be an increase in theta function in the hippocampus, with greater peak theta power in the Ts65Dn animals, and no difference in overall theta-band power. However, closer examination shows that theta is abnormal in Ts65Dn animals and occurs at higher frequencies than control. This abnormal theta frequency occurs alongside critical changes in the relationship of theta to the activity of single units and changes in the phasic relationships between theta and gamma that help explain how apparent increases in theta dominance are correlated with decreases in memory function.

Place cells in the hippocampus typically precess through phases of theta as animals traverse the active region of the cell^32–35,49,53–55^. In the present study, place cells recorded from non-trisomic control animals occur at all phases of theta. However, in Ts65Dn animals, these spikes occur at much more rigid phasic relationships to theta. Segregation of firing of single units on the theta wave is thought to gate information flow into epochs of encoding and retrieval based on the synchronicity between regions at specific points on the hippocampal theta cycle ^21,22^ and normal precession Likewise, in the 2N control animals in the present study, we see a recapitulation of the segregation of fast and slow gamma onto different phases of the theta cycle. Interestingly, there is increased coherence between theta and gamma overall in the Ts65Dn animals, again seemingly in contradiction to the known learning deficits show in Ts65Dn. However, the phasic relationship between theta and fast gamma is disturbed in these animals. While slow gamma occurred at similar theta phases to control animals, fast gamma was as likely to occur at the peak of theta as at the trough. This failure to segregate information switching from CA3 to MEC (and from retrospective to prospective coding) may underlie the learning deficits seen in both Ts65Dn animals, and humans with DS.

Exactly why the phasic segregation of hippocampal activity on the theta wave is abnormal in Ts65Dn animals is unknown, but the observation that GABA antagonists can correct some of the cognitive consequences of DS, and emerging evidence that theta within the hippocampus is controlled through GABAergic interneurons^56^ suggest interneurons synapsing onto CA1 pyramidal cells may drive this abnormal phase locking. PV+ interneurons are tightly phase-locked to the peak of theta, while SOM+ interneurons show a wider range of phasic relationships with theta^57^. One possibility for the abnormal phasic relationships seen in Ts65Dn animals is excessive interneuronal contact with place cells, either through greater interneuronal number, or a larger number of synapses. Although modelling work suggests that theta within CA1 can be produced without direct septal input, septal input is critically involved in regulating phasic relationships with theta^58^. An imbalance in the ratio of inhibitory GABAergic to excitatory cholinergic/glutamatergic projections from MS to HPC in Ts65Dn animals may be sufficient to perturb phasic relationships with theta within the hippocampus.

Given recent data suggesting that artificially prolonged sharp-wave ripples correlate with better spatial memory^43^, we were initially puzzled that Ts65Dn animals displayed ripples that were dramatically longer, on average, than 2N controls. However, a critical point is that longer-duration intrinsically generated sharp-wave ripples are correlated with increased demands on memory; that is, longer duration ripples indicate that the memory system is working harder^43^. In these experiments, animals had no special task demands, and the sharp-wave ripples of 2N control animals were predictably short. The longer duration ripples of Ts65Dn animals may illustrate their relatively worse memory performance generally; indeed environmental novelty is associated with longer duration ripples^43^. This may suggest that the environment was perpetually novel to the Ts65Dn animals, while the 2N control animals rapidly learned the spatial configuration of the recording chamber.

In these experiments, we find electrophysiological evidence of perturbation in hippocampal function in the Ts65Dn mouse model of DS that are consistent with the learning and memory deficits seen in this model. As in many other perturbation experiments, the gross appearance of place cells in these animals is normal, but the central changes are in the phasic relationship between place cells and theta rhythm, and between theta and other types of oscillation in LFP. This constellation of changes matches the hypothesis that the critical deficit in DS is output from medial septum; we find that changes in entrainment to theta rhythm are common to almost all the differences between Ts65Dn and control animals in these experiments.

## Acknowledgements

This project was supported by the LuMind Foundation. We would like to thank Dr. Elsa Pittaras for the breeding and supply of animals.

## Materials and Methods

### Animals and Surgery

All experimental procedures were carried out in accordance with and were approved by the Institutional Animal Care and Use Committee at Stanford University School of Medicine. Mice trisomic for the distal portion of chromosome 16 (Ts65Dn, n = 4) and their non-trisomic (2N Control, n = 4) littermates were used for single unit recordings. These animals were between 3 and 9 months of age at time of surgery, and weighed between 25 and 35 g. Animals in each cohort were from the same litter, and so were age-matched. Animals were deeply anaesthetised using isofluorane anaesthesia (1.5 – 2% in oxygen with flow rate 1200 – 1500 ml/min), and were given a subcutaneous injection of buprenorphine (0.03 mg). Animals were implanted with 16-channel tetrode carrying microdrives into the right hemisphere. The tetrodes themselves each consisted of four 17 µm Platinum/Iridium Polyimide-coated electrodes twisted together. The electrodes were electroplated using Platinum Black (Neuralynx) until their impedances were similar and were in the region 200 – 250 kΩ. Coordinates for the electrodes were −1.8 mm posterior to bregma, and 1.6mm lateral from the midline. Electrodes were implanted to an initial depth of 900-1000 µm, which typically placed them just dorsal to the hippocampal pyramidal cell layer. Animals were between 4 and 12 months old during recordings. Recordings were made during the animal’s daylight cycle.

### Statistics

All statistics were two-sided. A Lillefors test was used to determine whether the data were parametric. In the case of nonparametric data, either a signed-rank (paired) or rank-sum (unpaired) test was used. Differences between non-circular distributions were determined using a two-sample Kruskall-Wallis test. Determination of circular clustering was done via Rao’s spacing test. A nonparametric multisample test for equal circular medians was performed as in Fisher & Fisher (1993)^59^.

### Data Acquisition

Animals were screened for single units while they explored a 30 x 30 cm environment made of black polycarbonate. The walls of this enclosure were made unique through affixing a sheet of white paper to one wall, and alternating black and white diagonal stripes on the opposite wall. The orthogonal walls remained black. Local field potentials were co-recorded with single units, and were lowpass filtered from 0 – 500 Hz with a notch filter at 60 Hz. This channel was acquired at low resolution (250 Hz) and high resolution (4.8 kHz). The high resolution trace was used for all spectral analyses.

### Identification of firing fields

Spikes were converted into a binned ratemap, which summed the number of spikes in each 2 cm x 2 cm bin. This ratemap was then adaptively smoothed as in Skaggs et al. (1996)^34^. Regions of maximal activity were determined using the imextendedmax function in MATLAB v2019b. This function uses an 8-way connected region to determine the groups of bins that represent the maximal regions of firing. Each of these connected maxima neighbourhoods was defined as a field, and they were considered separate fields if the maximal regions shared no adjacent bins.

### Place cell classification

Cells were initially considered to be place cells based on spatial spike shuffling criteria for spatial sparsity, information content, and spatial selectivity. This shuffling procedure moved each spike time a random amount in both the x and y direction between the minimum and maximum x and y values explored in each recording session. We initially considered cells that were beyond the 99^th^ percentile of these values and met the other cell inclusion criteria (more than 100 spikes, more than 70% environmental coverage). In addition, cells had to have at least one identified firing field (see firing field classification), and fields had to cover at least 5% but not more than 60% of the environment. This classification had the effect of removing putative interneurons from the analyses, which tended to have non-discriminatory “fields” covering most of an environment.

### Pass analysis

This analysis followed Climer et al (2013)^49^. Place fields were identified using a “field index” computed using the bsxfun function in MATLAB, this provided a normalized global estimate over the map of how “in field” each position was (from 0 to 1). The centroids of fields were extracted using the regionprops function in MATLAB, and then a direction-invariant pass index through the fields was calculated using Hilbert transform hibert in MATLAB, which classified each pass through fields from −1 (start of pass), 0 (center of pass), to 1 (end of pass). The linearisation of passes in this way, allowed a linear-circular regression of spikes through the linear pass space and the circular LFP phase space. To eliminate spurious fits, the fits were constrained to have a negative slope between 0 and −2.

### Spectral analysis of theta rhythms

High-resolution LFP was bandpass filtered between 5 and 12 Hz, and then Fourier transformed. Instantaneous frequency and phase were determined by a Hilbert transform. To obtain estimates of the slope and the intercept of the relationship between running speed and frequency, a linear regression was performed on the running speed/frequency data. For the determination of spectral power in the theta band, a power spectrum of the high-resolution LFP trace was made using the pspectrum function in MATLAB 2019b with a frequency resolution of 0.5 Hz. Since spectral power can vary from animal to animal and depend on the duration of recording and the depth of the electrodes within the hippocampus, power was normalized within each recording such that the integral of the power between 3 and 30Hz was 1 (as in Russell et.al (2006)^60^).

### Spectral analysis of gamma rhythms and cross-frequency analysis

Analysis of gamma rhythms and cross-frequency coupling was modified from a previous technique^38^. High-resolution (4.8 kHz) LFP was transformed using a Morlet’s wavelet technique to determine the time-varying power in 2 Hz bins from 2-160 Hz. Spectral coherence between the original signal and the time-varying power was estimated using a magnitude squared function which consisted of a Hanning window FFT using 50% overlap. Individual gamma episodes were detected using the average time-varying power in the slow (25-50Hz) and fast (65-140Hz) bands. Gamma was detected when LFP epochs displayed power more than 2 standard deviations above the average power in each band across the recording. Gamma was then windowed as in Colgin et. al. (2009)^38^. The relationship of these gamma epochs to the theta phase was determined using a Hilbert transform to derive instantaneous phase of theta, and this was then referenced to the time of maximum power for each gamma epoch. Duplicated gamma episodes caused by overlapping time windows were removed by discarding gamma episodes with equal maxima, and requiring that each subtype of gamma be temporally separated by at least 100 ms.

### Spectral analysis of sharp-waves

High-resolution (4.8kHz) LFP was filtered using a Morlet’s wavelet centred on 200Hz with a bandwidth of 50Hz, giving a filter band of 150-250Hz. Detection of sharp waves on the resultant filtered signal was determined similarly to what was done previously^61^, epochs of filtered LFP were determined to be sharp waves if they were more than 3 standard deviations above the amplitude of the filtered signal, and that this magnitude persisted for 15ms or longer.

### Histology and determination of electrode position

At the conclusion of experiments animals were deeply anaesthetised with isoflurane, and then were injected with an overdose of sodium pentobarbitone intraperitoneally. Animals were then perfused transcardially with 0.9% phosphate buffered saline, followed by 9% formalin. Animals were decapitated, and their brains removed and placed into a cylindrical container holding a solution of 9% formalin for 24-48h. Brains were then transferred to a 30% sucrose solution, where they were kept until they sunk from the top of the container to the bottom. Brains were removed from solution, quickly frozen, and then sectioned coronally into slices of 40 micron thickness. Slices were mounted, and then Nissl stained to visualise cell bodies and make the electrode positions apparent. The track made by the tetrodes and its end position was determined through microscopic examination of the slides, and the depth at which each cell was recorded was determined by back-calculation from this final location.

## References

1. Moraes, M. E. L. de, Tanaka, J. L. O. Moraes, L. C. de, Filho, E. M. & Castilho, J. C. de M. Skeletal age of individuals with Down syndrome. Special Care Dent 28, 101–106 (2008).

2. Angelopoulou, N. et al. Bone Mineral Density and Muscle Strength in Young Men with Mental Retardation (With and Without Down Syndrome). Calcified Tissue Int 66, 176–180 (2000).

3. Marino, B. et al. Atrioventricular Canal in Down Syndrome: Prevalence of Associated Cardiac Malformations Compared With Patients Without Down Syndrome. Am J Dis Child 144, 1120–1122 (1990).

4. Chaoui, R. et al. Aberrant right subclavian artery as a new cardiac sign in second- and third-trimester fetuses with Down syndrome. Am J Obstet Gynecol 192, 257–263 (2005).

5. Yamaki, S., Horiuchi, T. & Sekino, Y. Quantitative analysis of pulmonary vascular disease in simple cardiac anomalies with the down syndrome. Am J Cardiol 51, 1502–1506 (1983).

6. Rowe, J., Lavender, A. & Turk, V. Cognitive executive function in Down’s syndrome. Brit J Clin Psychol 45, 5–17 (2006).

7. Lanfranchi, S., Jerman, O., Pont, E. D., Alberti, A. & Vianello, R. Executive function in adolescents with Down Syndrome. J Intell Disabil Res 54, 308–319 (2017).

8. Simon, E. W., Rappaport, D. A. & Agriesti, M. Memory performance in adults with Down syndrome. Australia New Zealand J Dev Disabil 20, 113–125 (2009).

9. Lott, I. T. & Dierssen, M. Cognitive deficits and associated neurological complications in individuals with Down’s syndrome. Lancet Neurology 9, 623–633 (2010).

10. Pennington, B. F., Moon, J., Edgin, J., Stedron, J. & Nadel, L. The Neuropsychology of Down Syndrome: Evidence for Hippocampal Dysfunction. Child Dev 74, 75–93 (2003).

11. Kelley, C. M. et al. Maternal choline supplementation differentially alters the basal forebrain cholinergic system of young-adult Ts65Dn and disomic mice. J Comp Neurol 522, 1390–1410 (2014).

12. Kelley, C. M. et al. Sex differences in the cholinergic basal forebrain in the Ts65Dn mouse model of Down syndrome and Alzheimer’s disease. Brain Pathology Zurich Switz 24, 33–44 (2013).

13. Vandecasteele, M. et al. Optogenetic activation of septal cholinergic neurons suppresses sharp wave ripples and enhances theta oscillations in the hippocampus. Proc National Acad Sci 111, 13535–13540 (2014).

14. Allen, C. N. & Crawford, I. L. GABAergic agents on the medial septal nucleus affect hippocampal theta rhythm and acetylcholine utilization. Brain Res 322, 261–267 (1984).

15. Brazhnik, E. S. & Vinogradova, O. S. Control of the neuronal rhythmic bursts in the septal pacemaker of theta-rhythm: Effects of anaesthetic and anticholinergic drugs. Brain Res 380, 94–106 (1986).

16. Vinogradova, O. Expression, control, and probable functional significance of the neuronal theta-rhythm. (1995).

17. Brazhnik, E. S., Vinogradova, O. S. & Karanov, A. M. Frequency modulation of neuronal theta-bursts in rabbit’s septum by low-frequency repetitive stimulation of the afferent pathways. Neuroscience 14, 501–508 (1985).

18. Newman, E. L., Gillet, S. N., Climer, J. R. & Hasselmo, M. E. Cholinergic blockade reduces theta-gamma phase amplitude coupling and speed modulation of theta frequency consistent with behavioral effects on encoding. J Neurosci Official J Soc Neurosci 33, 19635–46 (2013).

19. Newman, E. L. et al. Precise spike timing dynamics of hippocampal place cell activity sensitive to cholinergic disruption. Hippocampus 27, 1069–1082 (2017).

20. Schomburg, E. W. et al. Theta Phase Segregation of Input-Specific Gamma Patterns in Entorhinal-Hippocampal Networks. Neuron 84, 470–85 (2014).

21. Hasselmo, M. E., Bodelón, C. & Wyble, B. P. A proposed function for hippocampal theta rhythm: separate phases of encoding and retrieval enhance reversal of prior learning. Neural computation 14, 793–817 (2002).

22. Manns, J. R., Zilli, E. A., Ong, K. C., Hasselmo, M. E. & Eichenbaum, H. Hippocampal CA1 spiking during encoding and retrieval: Relation to theta phase. Neurobiology of Learning and Memory 87, 9–20 (2007).

23. Hasselmo, M. E. & Stern, C. E. Theta rhythm and the encoding and retrieval of space and time. Neuroimage 85, 656–666 (2014).

24. Judge, S. J. & Hasselmo, M. E. Theta Rhythmic Stimulation of Stratum Lacunosum-Moleculare in Rat Hippocampus Contributes to Associative LTP at a Phase Offset in Stratum Radiatum. Journal of Neurophysiology 92, 1615–1624 (2004).

25. Hasselmo, M. E. What is the function of hippocampal theta rhythm?—Linking behavioral data to phasic properties of field potential and unit recording data. Hippocampus 15, 936–949 (2005).

26. Boyce, R., Glasgow, S. D., Williams, S. & Adamantidis, A. Causal evidence for the role of REM sleep theta rhythm in contextual memory consolidation. Sci New York N Y 352, 812–6 (2016).

27. Mojabi, F. S. et al. GABAergic hyperinnervation of dentate granule cells in the Ts65Dn mouse model of down syndrome: Exploring the role of App. Hippocampus 26, 1641–1654 (2016).

28. Kleschevnikov, A. M. et al. Increased efficiency of the GABAA and GABAB receptor-mediated neurotransmission in the Ts65Dn mouse model of Down syndrome. Neurobiol Dis 45, 683–91 (2011).

29. Hangya, B., Borhegyi, Z., Szilágyi, N., Freund, T. F. & Varga, V. GABAergic Neurons of the Medial Septum Lead the Hippocampal Network during Theta Activity. J Neurosci 29, 8094–8102 (2009).

30. Gulyás, A. I., Görcs, T. J. & Freund, T. F. Innervation of different peptide-containing neurons in the hippocampus by gabaergic septal afferents. Neuroscience 37, 31–44 (1990).

31. Costa, E., Panula, P., Thompson, H. K. & Cheney, D. L. The transsynaptic regulation of the septal-hippocampal cholinergic neurons. Life Sci 32, 165–179 (1983).

32. Maurer, A. P., Cowen, S. L., Burke, S. N., Barnes, C. A. & McNaughton, B. L. Organization of hippocampal cell assemblies based on theta phase precession. Hippocampus 16, 785–794 (2006).

33. Bose, A. & Recce, M. Phase precession and phase-locking of hippocampal pyramidal cells. Hippocampus 11, 204–215 (2001).

34. Skaggs, W. E., McNaughton, B. L., Wilson, M. A. & Barnes, C. A. Theta phase precession in hippocampal neuronal populations and the compression of temporal sequences. Hippocampus 6, 149–172 (1996).

35. Tsodyks, M. V., Skaggs, W. E., Sejnowski, T. J. & McNaughton, B. L. Population dynamics and theta rhythm phase precession of hippocampal place cell firing: A spiking neuron model. Hippocampus 6, 271–280 (1996).

36. Kunec, S., Hasselmo, M. E. & Kopell, N. Encoding and Retrieval in the CA3 Region of the Hippocampus: A Model of Theta-Phase Separation. Journal of Neurophysiology 94, 70–82 (2005).

37. Bieri, K. W., Bobbitt, K. N. & Colgin, L. L. Slow and Fast Gamma Rhythms Coordinate Different Spatial Coding Modes in Hippocampal Place Cells. Neuron 82, 670–681 (2014).

38. Colgin, L. et al. Frequency of gamma oscillations routes flow of information in the hippocampus. Nature 462, 353 (2009).

39. Fernández-Ruiz, A. et al. Entorhinal-CA3 Dual-Input Control of Spike Timing in the Hippocampus by Theta-Gamma Coupling. Neuron 93, 1213-1226.e5 (2017).

40. Pfeiffer, B. E. & Foster, D. J. Hippocampal place cell sequences depict future paths to remembered goals. Nature 497, 74 (2013).

41. Foster, D. J. & Wilson, M. A. Reverse replay of behavioural sequences in hippocampal place cells during the awake state. Nature 440, 680–683 (2006).

42. Jadhav, S. P., Kemere, C., German, W. P. & Frank, L. M. Awake Hippocampal Sharp-Wave Ripples Support Spatial Memory. Science 336, 1454–1458 (2012).

43. Fernández-Ruiz, A. et al. Long-duration hippocampal sharp wave ripples improve memory. Science 364, 1082–1086 (2019).

44. Fernandez, F. et al. Pharmacotherapy for cognitive impairment in a mouse model of Down syndrome. Nat Neurosci 10, 411–413 (2007).

45. Colas, D., Chuluun, B., Garner, C. C. & Heller, H. C. Short-term treatment with flumazenil restores long-term object memory in a mouse model of Down syndrome. Neurobiol Learn Mem 140, 11–16 (2017).

46. Colas, D. et al. Short-term treatment with the GABAA receptor antagonist pentylenetetrazole produces a sustained pro-cognitive benefit in a mouse model of Down’s syndrome. Brit J Pharmacol 169, 963–973 (2013).

47. Chang, P. et al. Altered Hippocampal-Prefrontal Neural Dynamics in Mouse Models of Down Syndrome. Cell Reports 30, 1152-1163.e4 (2020).

48. McFarland, W. L., Teitelbaum, H. & Hedges, E. K. Relationship between hippocampal theta activity and running speed in the rat. Journal of Comparative and Physiological Psychology 88, 324 (1975).

49. Climer, J. R., Newman, E. L. & Hasselmo, M. E. Phase coding by grid cells in unconstrained environments: two-dimensional phase precession. European J Neurosci 38, 2526–41 (2013).

50. Climer, J. R., DiTullio, R., Newman, E. L., Hasselmo, M. E. & Eden, U. T. Examination of rhythmicity of extracellularly recorded neurons in the entorhinal cortex. Hippocampus 25, 460–473 (2015).

51. Carr, M. F., Karlsson, M. P. & Frank, L. M. Transient Slow Gamma Synchrony Underlies Hippocampal Memory Replay. Neuron 75, 700–13 (2012).

52. Reeves, R. H. et al. A mouse model for Down syndrome exhibits learning and behaviour deficits. Nat Genet 11, 177–184 (1995).

53. Maurer, A. P. & McNaughton, B. L. Network and intrinsic cellular mechanisms underlying theta phase precession of hippocampal neurons. Trends in Neurosciences 30, 325–333 (2007).

54. Jaramillo, J. & Kempter, R. Phase precession: a neural code underlying episodic memory? Curr Opin Neurobiol 43, 130–138 (2017).

55. O’Keefe, J. & Recce, M. L. Phase relationship between hippocampal place units and the EEG theta rhythm. Hippocampus 3, 317–330 (1993).

56. Bezaire, M. J., Raikov, I., Burk, K., Vyas, D. & Soltesz, I. Interneuronal mechanisms of hippocampal theta oscillations in a full-scale model of the rodent CA1 circuit. Elife 5, e18566 (2016).

57. Huh, C. Y. L. et al. Excitatory Inputs Determine Phase-Locking Strength and Spike-Timing of CA1 Stratum Oriens/Alveus Parvalbumin and Somatostatin Interneurons during Intrinsically Generated Hippocampal Theta Rhythm. J Neurosci Official J Soc Neurosci 36, 6605–22 (2016).

58. Mysin, I. E., Kitchigina, V. F. & Kazanovich, Y. B. Phase relations of theta oscillations in a computer model of the hippocampal CA1 field: Key role of Schaffer collaterals. Neural Networks 116, 119–138 (2019).

59. Fisher, N. I. & Fisher, N. Statistical Analysis of Circular Data. xv–xviii (1993) doi:10.1017/cbo9780511564345.002.

60. Russell, N. A., Horii, A., Smith, P. F., Darlington, C. L. & Bilkey, D. K. Lesions of the Vestibular System Disrupt Hippocampal Theta Rhythm in the Rat. Journal of Neurophysiology 96, 4–14 (2006).

61. Gillespie, A. K. et al. Apolipoprotein E4 Causes Age-Dependent Disruption of Slow Gamma Oscillations during Hippocampal Sharp-Wave Ripples. Neuron 90, 740–751 (2016).

